# Improved mRNA electroporation into immature oocytes to examine protein dynamics during development

**DOI:** 10.1101/2023.12.07.570343

**Authors:** Yuhkoh Satouh, Emiko Suzuki, Keisuke Sasaki, Ken Sato

## Abstract

One of the major cause of oocyte quality deterioration along with aging, chromosome segregation abnormalities occur mainly during meiosis I. However, currently, there is a technical limitation in the introduction of mRNA into premature oocytes without impairing embryonic developmental ability. In this study, we established a low-invasive electroporation (EP) method to introduce mRNA into pre-ovulatory, germinal vesicle (GV) mouse oocytes in an easier manner than the traditional microinjection method. The EP method with an optimized impedance value resulted in the efficient introduction of mRNAs encoding enhanced green fluorescent protein (EGFP) into the GV oocytes surrounded by cumulus cells at a survival rate of 95.0%. Furthermore, the introduction of histone H2B-EGFP mRNA into the GV oocytes labeled most of the oocytes without affecting the blastocyst development rate, indicating the feasibility of the visualization of oocyte chromosomal dynamics that enable us to assay chromosomal integrity in oocyte maturation and cell count in embryonic development. The establishment of this EP method offers extensive assays to select pre-implantation embryos and enables the surveying of essential factors for mammalian oocyte quality determination.

**Summary blurb:** This study introduces a low-invasive electroporation method with high survival rate and developmental ability, offering a potential breakthrough in preimplantation embryo treatment and assessment.

## Introduction

Currently, the major cause of female infertility is the deteriorated quality of oocytes which is potentially manifested by the increased childbearing age of females (Secomandi et al., 2022). Although the biological basis underlying such changes in oocyte quality is not fully understood, the aberrations of oocyte chromosomes mainly caused by segregation failure are obviously correlated with the reduced developmental ability (Vazquez-Diez and FitzHarris, 2018). However, treatment of such chromosomal aberrations has been faced with two major difficulties. One is concerns that interventions in the oocyte chromosomes themselves include risks that may cause genetic manipulation, while another is technical difficulties in introducing exogenous factors to fluctuate the oocyte quality or monitor the oocyte chromosomes, in a low- or non-invasive manner.

In mammals, a fully-grown primary oocyte, which has the obvious germinal vesicle (GV), arrests at the first meiotic division (MI) in the ovarian follicle and releases the first polar body with chromosomes during or after ovulation, a transition from the ovary to the oviduct. The ovulated oocytes of mice and humans arrest their cell cycles and cellular dynamics again in the metaphase of the second meiotic division (MII), awaiting the sperm to be fertilized in the oviduct. Unfertilized oocytes are thus called MII oocytes (Bakloushinskaya, 2022; Conti and Franciosi, 2018; Zielinska et al., 2019). The transition from a GV to an MII oocyte is called oocyte maturation and can be induced in vitro with an in vitro maturation (IVM) treatment. Recent studies that tracked the incidents of such chromosomal aberrations have shown that the errors in chromosomal segregation in aged mammalian oocytes take place mainly at the time of the first polar body release, during the oocyte maturation stage (Sakakibara et al., 2015; Thomas et al., 2021). These facts suggest that the most possible and effective solution to prevent chromosome aberration due to aging is an intervention at the pre-ovulatory stage such as the GV stage, in which the harvested oocytes need to be matured in vitro.

In recent years, owing in part to the great advances in in vitro oogenesis technologies, MII oocytes that can support embryonic development at a higher rate than before are obtained by IVM using mouse GV oocytes or oocytes from the earlier follicle stages (Kohama et al., 2022; Morohaku et al., 2017; Morohaku et al., 2016). Moreover, lower-invasive types of monitoring methods for the visualization of chromosomal integrities have been established, including confocal or lattice light sheet microscopic time-lapse observations using fluorescent markers (Hatano et al., 2022; Qin et al., 2017; Yamagata and Ueda, 2013). However, technical limitations have existed for introducing solutions containing proteins, mRNA, siRNA, morpholino, etc. into the pre-ovulatory oocytes. The most used method to introduce these solutions into the GV oocytes is microinjection using a glass needle. The limitations with the microinjection are not only due to the technical difficulties of executing this method because of the sensitive and fragile nature of the GV oocytes but also because of the requirement of needing to remove the cumulus cells that are tightly attached to the zona pellucida (ZP) to visualize an ooplasm boundary. The cumulus cells play an important role in supporting oocyte maturation and fertilization, and their removal has been reported to reduce the incidence of embryos derived from oocytes following IVM (Gilchrist and Smitz, 2023). Thus, the low-invasive introduction of solutions and conservation of the cumulus cells are required when establishing effective methods to intervene in GV oocytes. Recently, electroporation (EP) based methods have been applied broadly to introduce solutions into oocytes or embryos in a low-invasive manner, such as the TAKE method in which solutions containing the Cas9 protein and guide RNAs are introduced into mouse/rat zygotes for gene editing (Kaneko and Mashimo, 2015; Kaneko et al., 2014). Therefore, this study aimed to establish a method to introduce mRNA encoding marker proteins into GV oocytes, as easily as possible, noninvasively, and with as many cumulus cells preserved as possible, using EP.

## Results

### Design of a low-invasive EP method into GV oocytes

GV oocytes collected in vivo are diverse in that some oocytes have more, less, or no cumulus; oocytes enclosed with more cumulus cells are shown to present better developmental abilities in IVM (Schroeder and Eppig, 1984; Vanderhyden and Armstrong, 1989). In the present study, we designated those with more than 90% of the ZP surface exposed as cumulus (-), those with a portion exposed or covered by a single layer of cumulus cells as cumulus (+), and those with no ZP exposure and enclosed with multiple layers of cumulus cells as cumulus (++). For the C57BL/6NCrSlc mice with which this experiment was conducted, cumulus (-) was collected at 27.1%, cumulus (+) at 22.8%, and cumulus (++) at 50.1% (total 403 oocytes: **Fig 1A**).

**Figure 1.**
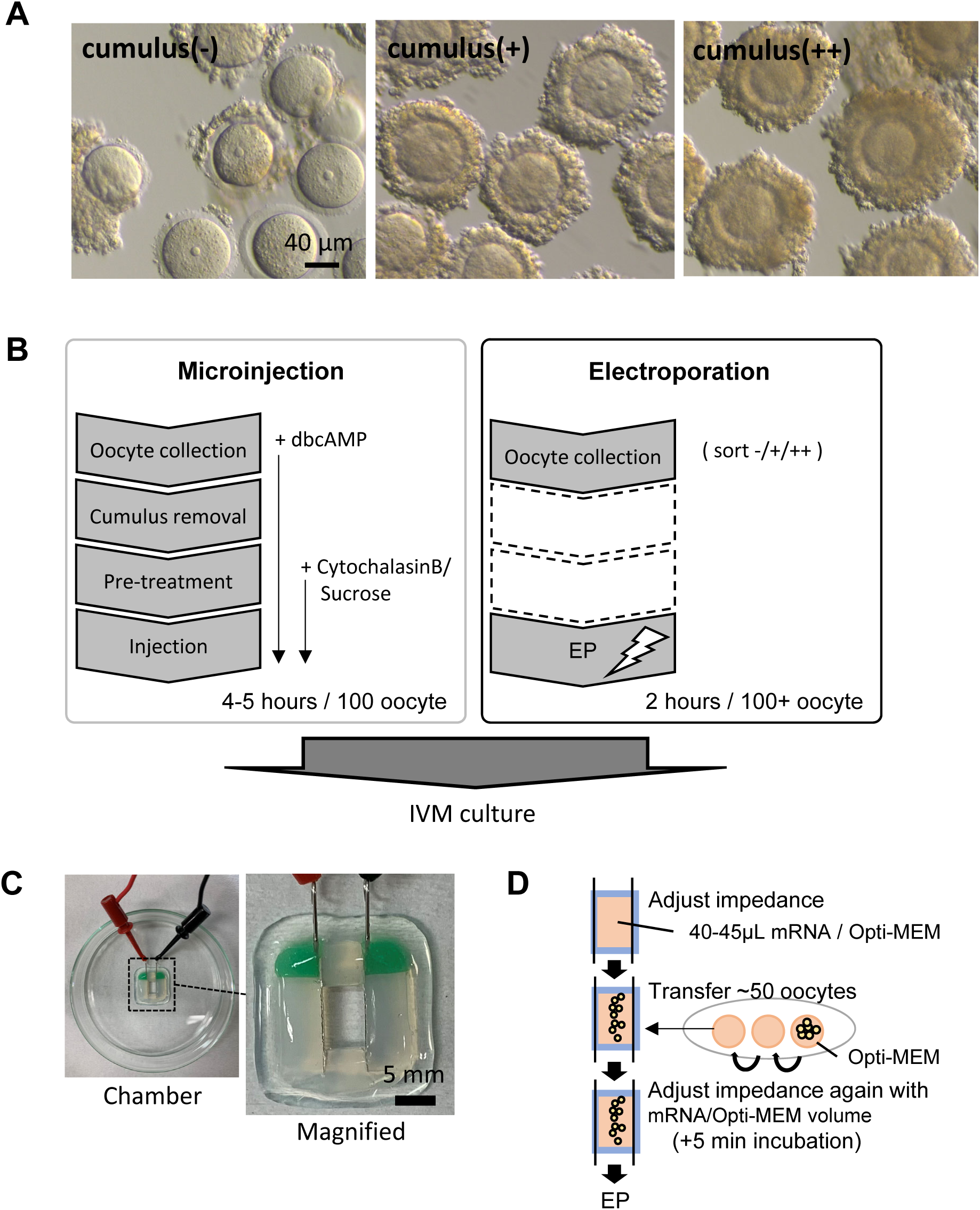
Experimental design for EP of GV mouse oocytes. A) GV oocytes with more than 90% of the ZP surface exposed were designated as cumulus (-), those partially exposed or covered by a single layer of cumulus were designated as cumulus (+), and those with no exposure and covered by multiple layers of cumulus were designated as (++). B) In the conventional microinjection experiment, oocytes were collected in a medium supplemented with dbcAMP. After the cumulus removal with a micromanipulator, CCB and 0.25 M Sucrose were additively supplemented to avoid fatal damage due to mechanical stress. In the EP experiment, oocytes were immediately subjected to EP treatment after being sorted into the cumulus (-), (+), or (++) groups. C) GV oocytes were placed in the EP chamber. The same number of chambers were prepared for all the experimental batches. D) Procedure for low-invasive EP. Impedance was first set to the target value and then adjusted again after placing the oocytes (just before electroporation).

The microinjection was carried out following conventional methods; due to the long process of the method (3+ h), dibutyryl-cyclic AMP (dbcAMP), which prevents GV breakdown (GVBD) by arresting the cell cycle, was supplemented, and the cumulus was removed by mechanical aspiration with a micromanipulator as shown in **SFig 1**. To avoid damage to the plasma membrane of the very fragile GV oocyte by physical contact with the needle, the cytoskeleton underneath the plasma membrane was depolymerized by supplementation with Cytochalasin B (CCB), and injections were performed in the presence of 0.25 M Sucrose to shrink and give slack for needle insertion. However, for EP, after the classification of the described GV oocytes according to the number of cumulus cells as above, the cells were immediately transferred to the EP treatment (**Fig 1B**).

Since the electrical impedance of the EP chamber is a determinant of survival after EP, i.e., invasiveness, the impedance values were adjusted to around 500 Ω in the case of mammalian embryos (Yoshimatsu et al., 2019), and 520 Ω was adopted as the initial setting value in the TAKE method. To explore lower-invasive conditions, we additively adopted 590 Ω as a potentially less-invasive condition, and the impedance value of 520 Ω or 590 Ω was accurately set by adjusting the solution volume just prior to EP (**Figs 1C, D**).

### EP into GV oocytes allows low-invasive introduction of mRNA

Under these conditions described above, we introduced mRNA encoding EGFP, whose fluorescence could be detected uniformly in the cytosol, to quantify the efficiency of mRNA introduction and the invasiveness of the EP operation by oocyte survival. To make the comparison as simple as possible, the same concentration (200 ng/μL) was used both for the external solution of EP and infusion of the microinjection. The survival rate for each treatment after a 17-h-incubation of IVM was not significantly different but that of cumulus (++) for 590 Ω was the highest among the treated groups (**Fig 2A**). In the no-treatment group, cumulus (++) demonstrated a significantly better MII rate than cumulus (-) and cumulus (+), supporting previous reports (Schroeder and Eppig, 1984; Vanderhyden and Armstrong, 1989). However, in EP 520 Ω, but not in EP 590Ω, the advantage of cumulus (++) was lost, and it was not significantly different (**Fig 2B**). In comparison to the cumulus (++) batches and the injected oocytes, the injection treatment showed significantly reduced MII rates, while EP did not, against no-treatment cumulus (++). In particular, the cumulus (++) of 590 Ω showed a P-value of 0.997 versus that of no treatment, indicating a very low invasiveness result (**Fig 2B**).

**Figure 2.**
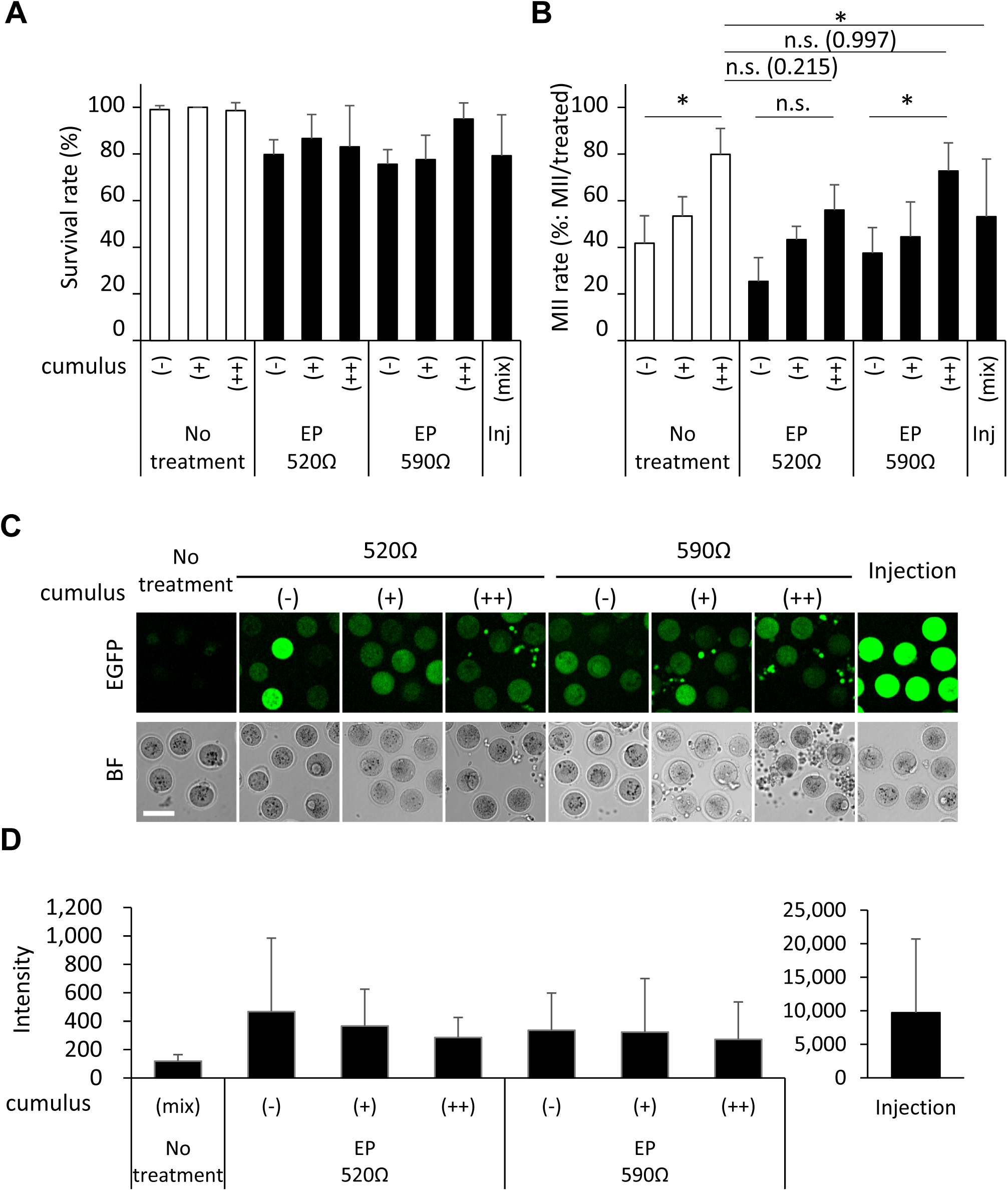
High impedance EP to cumulus-intact GV oocytes achieved low-invasive mRNA introduction. A) Survival rates indicate the averaged percentile of surviving oocytes per treated oocytes after IVM incubation. B) MII rates indicate the averaged percentile of MII oocytes per treated oocytes after IVM incubation. Significance was tested using one-way ANOVA (the P-values are indicated in parenthesis). Asterisks: significant; n.s.: not significant. C) Fluorescence captured with confocal microscopy. Bar: 100 μm. D) Fluorescent intensities captured in C). Error bars indicate the SD.

The fluorescence observation of the treated oocytes following IVM indicated that the introduction of EGFP into the cumulus (with strong bright spots) and into the oocytes (with dim and spread signals over the oocyte cytoplasm) was detected in all of the cumulus (+) and (++) EP batches (**SFig 2**). A more accurate evaluation of the amount of introduced mRNA after removal of the cumulus cells indicated that almost all of the treated oocytes gave a positive EGFP fluorescent signal; however, the no-treatment group showed only weak background signals (**Fig 2C**). As observed in **Fig 2D**, most of the oocytes in the treated batch showed higher intensities than the background level; however, the intensities varied greatly within the population, especially for 520 Ω (-). The amount of mRNA induction did not vary greatly depending on the Ω; however, the fluorescent intensities of the EP batches were about 1/20–1/40 that of the microinjection batch, in which the mRNA solution at the concentration of the external solution of the EP was directly introduced.

We then examined the introduction of EGFP into the MII oocytes. Almost all the MII oocytes collected in vivo from oviducts are encased in a cumulus cell layer; however, the expanded cumulus cell layer is easily removed by hyaluronidase treatment. Therefore, we tested in the same way for the 520 Ω and 590 Ω settings for the cumulus-intact (+) and removed (-), respectively. However, while microinjection of the MII stage oocytes (without any excess reagent treatment but with hyaluronidase) did not result in any invasiveness, the EPs were not only highly lethal but also frequently showed abnormal activation and fragmentation, such as the formation of pronuclear oocyte chromosomes (**SFig 3**).

### Chromosome dynamics during IVM can be visualized by introducing histone H2B-EGFP mRNA by EP

We then tested the feasibility of examining the oocyte chromosomal integrity under the EP condition of cumulus (++) GV oocytes, which gave the most non-invasive results, by introducing an mRNA coding histone H2B-EGFP, which is often used to assay chromosomal integrity in live imaging methods (Hatano et al., 2022; Yamagata et al., 2005). We examined using GV oocytes from a hybrid (B6N x DBA2 F1: BDF1), which are often used for developmental experiments of oocytes after IVM (Kohama et al., 2022; Morohaku et al., 2017; Morohaku et al., 2016).

When cumulus cells are present during IVM, the interior of the ZP cannot be clearly observed in bright field observation. However, histone H2B-EGFP fluorescence was observed within 4–5 hours after EP in almost all the EP oocytes, and the timings for the segregation of chromosomes from MI to MII and the release of the first polar bodies could be identified from the fluorescence signatures (**Fig 3A**). Both the lagging or the isolated chromosomes during the IVM time-lapse, and the chromosomal structure after IVM could be observed clearly (**Fig 3B, C**). Presence or absence of cumulus cells did not affect the detection of the chromosome, although the cumulus (++) oocytes tend to drift three dimensionally along with the expansion of cumulus cell layer during the oocyte maturation (**Supplementary Movies 1 and 2**).

**Figure 3.**
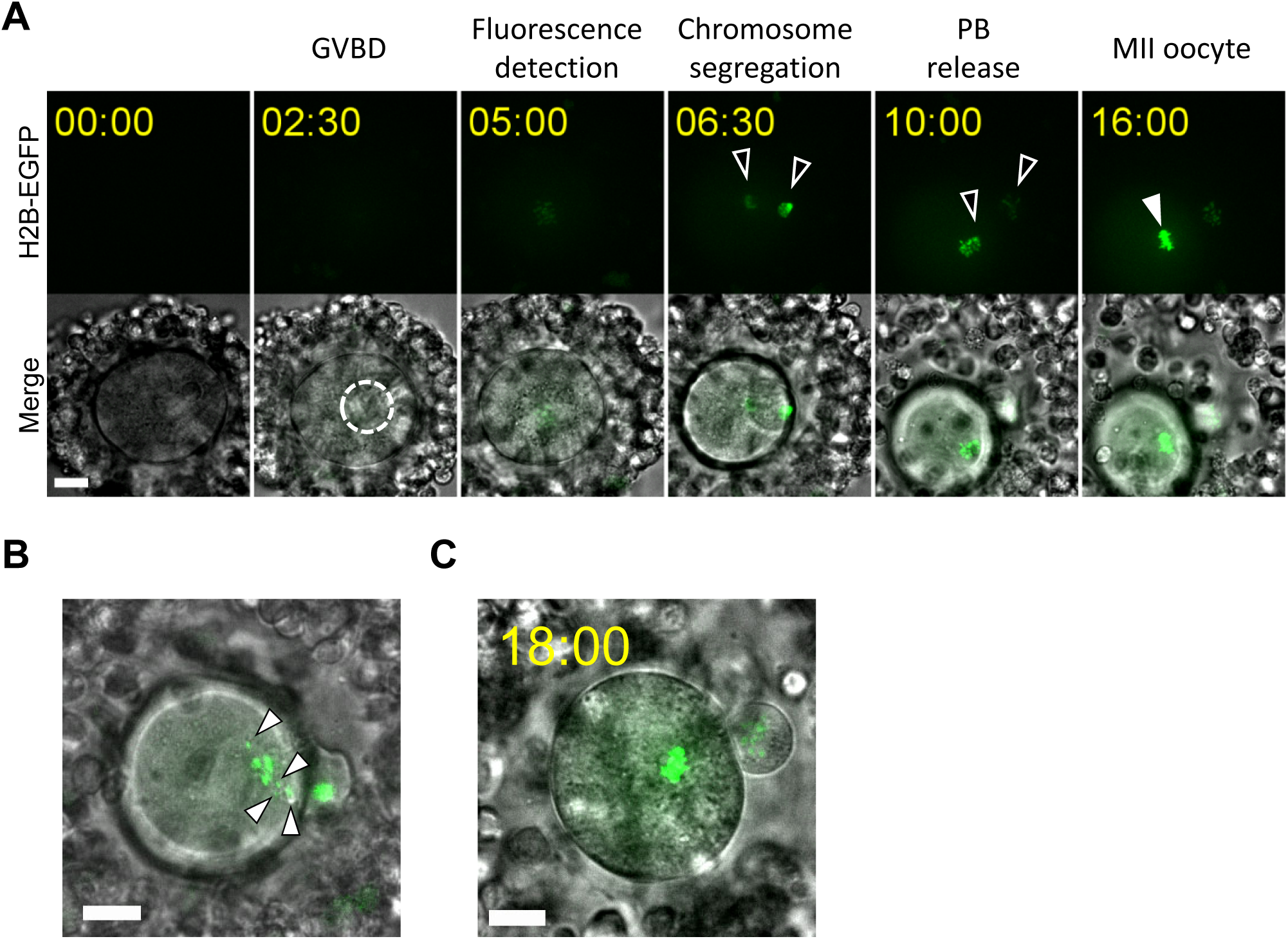
EP of the mRNA for histone H2B-EGFP specifically visualizes chromosomes during oocyte maturation. A) Time-lapse observation of IVM. Histone H2B-EGFP fluorescence observed on a laser confocal fluorescence microscope. Time stamps indicate the time after initiation of the image acquisition (just after EP and IVM culture). The black arrowheads indicate segregated chromosomes. The white arrowhead indicates the MII chromosome. B) Example of the isolated chromosomes observed in the IVM time-lapse (6 h elapsed). The arrowheads indicate isolated chromosomes. C) Chromosomes of an MII oocyte before in vitro fertilization (IVF). Bars: 20 μm. All the confocal images indicated are maximum intensity projections.

### Low-invasive EP into GV oocytes does not affect embryonic development

Finally, we tested whether this minimally invasive EP procedure affects embryonic development and whether the chromosome visualization enables us to track chromosomes during both fertilization and early embryonic development by performing in vitro fertilization and further embryonic culture experiments using embryos introduced with histone H2B-EGFP mRNA. Mouse embryos that have undergone IVM are known to require additional culture (120 hours instead of the usual 96 hours) to become blastocyst-stage embryos. Therefore, we analyzed the embryos based on the criterion of whether they had become pronuclear zygotes, 2-cell stage embryos, 4-cell stage embryos, morula embryos, blastocyst-stage embryos, and blastocyst-stage embryos at 6, 24, 48, 72, 96, and 120 hours, respectively.

The results showed that the blastocyst rate (to the number of MII oocytes) of the cumulus (++) GV oocytes without any treatment at 120 hpf was 31.7%, while that of the oocytes introduced with histone H2B-EGFP was 30.7%, which showed no significant reduction (**Fig 4A**). Although the number of experiment is small (N=1), 22.7 % (5/22 embryos) of embryos developed to blastocyst stage even when they are fertilized after a live-cell imaging during IVM. In addition, during the pronuclear stage (6 hpf), the pronuclei could be counted in all the embryos, which allowed for the evaluation of mono- or poly-spermy (**Fig 4B**). Furthermore, chromosomal fluorescence continued to be detected even after development to the blastocyst stage (120 hpf), enabling us to count the number of cells in the blastocyst embryos (**Fig 4C**).

**Figure 4.**
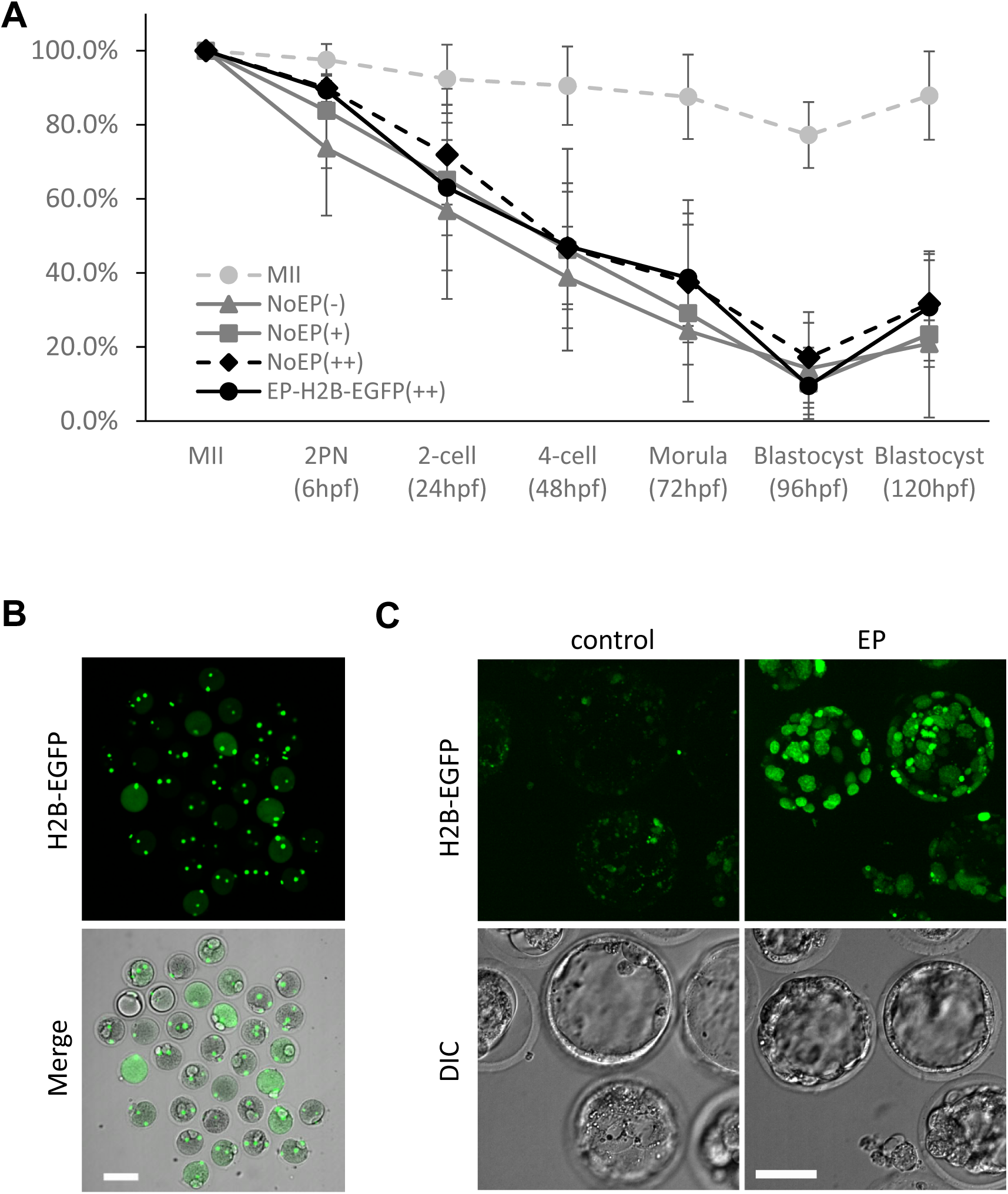
Cumulus-intact EP allows embryonic development and evaluation of pre-implantation embryos. A) Averaged rates of embryonic development after the EP to cumulus-intact GV oocytes of histone H2B-EGFP mRNA. Percentages to the number of MII oocyte for 2PN zygotes, 2-, 4-, 8-cell embryos, morula, blastocyst at each indicated hpf, respectively, are indicated. MII: harvested from the oviduct (matured in vivo). No EP: (-), (+), or (++) GV oocytes without EP. B) EGFP fluorescent of PN zygotes at 6 hpf observed using a laser confocal fluorescence microscope (maximum projection image is indicated). Bar: 100 μm. C) EGFP fluorescence of blastocyst embryos at 120 hpf observed using a laser confocal fluorescence microscope (maximum projection image is indicated). Control: (++) GV oocytes without EP. Bars: 40 μm.

## Discussion

In this study, we demonstrated that EP to cumulus-intact GV oocytes with optimized impedance can offer a minimally invasive introduction of mRNA and drive exogenous protein expression in mouse oocytes. When a fluorescent chromosome marker was introduced by this method, it was possible not only to observe the chromosomal integrity during IVM but also to examine chromosome dynamics during embryonic development and measure the number of cells. Although the EP condition was not applicable for MII oocytes, considering that the EP method has a generally lower technical barrier for introduction than the microinjection, this could be an advantage when manipulating the pre-ovulatory oocyte. The measurement of cell counts at the blastocyst stage is a promising index that affects post-transplantation performance and is expected to be applied to the selection of pre-implantation embryos.

We explored our EP conditions for the GV oocytes according to the TAKE method, which is currently widely used in genome editing methods using CRISPR/Cas9 (Kaneko et al., 2014), as a basic condition. As a result, we found that a relatively higher impedance value of 590 Ω allows for the lowest invasiveness and effective introduction of mRNAs into the GV oocytes enclosed with multiple layers of cumulus cells than the standard impedance value in the TAKE method and found that the MII oocyte is not suitable for EP with these settings. Yamamoto et al. recently reported a successful suppression of the H3f3a/b gene expression by introducing siRNA (−23 bp) into mouse GV oocytes using electroporation in a manner similar to our study (Yamamoto et al., 2023). In this study, we introduced a much longer size of mRNAs of approximately 1.2 kbp and succeeded in the efficient expression of fluorescent proteins in the oocytes and embryos. This improvement provides an important advancement in the search for genes to ameliorate oocyte quality and examine protein dynamics and functions during early development, by introducing the mRNAs of interest. At the same time, the feasibility of both the suppression and induction of protein expression is expected to have a synergistic effect in terms of future analysis of mammalian oocyte quality and as a tool to improve the quality of oocytes derived from patients.

We found that the introduction efficiency of mRNAs coding cytosolic EGFP or histone H2B-EGFP into oocytes was better than that into cumulus cells, suggesting a unique characteristic of oocytes in EP that originates from the relationship between surface area and volume. Considering the nature of electroporation, in that transportation from the extracellular environment takes place on the plasma membrane surface, the oocyte has a larger surface area than many other somatic cells (e.g., cumulus cells in this study) and is considered to have a higher mRNA transfection rate in terms of mRNA per cell. However, the number of chromosomes in both cells is at most twice as large (if the oocyte retained 4N), which may explain why the oocytes are positive at a higher frequency whereas the cumulus is not so fluorescently positive following the histone H2B-EGFP introduction (**Supplementary Movies 1 and 2**).

In this study, the blastocyst development rate of oocytes after EP, or no treatment, is approximately 30% equivalent to those of mouse embryos obtained via other IVM studies (Kohama et al., 2022; Morohaku et al., 2016), which indicates that the EP treatment developed here does not adversely affect the oocyte developmental ability. Furthermore, we interpreted the IVM oocytes’ developmental rate, allowing us to observe a significant promotive and suppressive effect on the incidence. Moreover, the blastocyst rate of the oocytes collected at MII (matured in vivo) was much higher (87.9% in this study) than those of the oocytes obtained via IVM; similarly, in vitro gametogenesis with more primitive germ cells or pluripotent stem cells are still subjected to a decreased oocyte quality (Hikabe et al., 2016; Morohaku et al., 2016). These facts indicate that in vitro steps for obtaining mammalian oocytes need to be improved, which in turn will enable us to explore promoting factors that can be commonly supplemented for in vitro generated oocytes by introducing some mRNAs or other reagents along with the method of this study.

Finally, further improvement of the EP technique and development of instrumentation would further reduce the invasiveness of the assay for pre-ovulatory oocytes and expand its range of application by increasing the amount of transduction. It is hoped that the results of this study will lead not only to predictions of oocyte quality but also to the development of IVM technology, and proactive interventions that will improve oocyte quality and avoid the effects of aging.

## Materials and Methods

### Animal experiments and strains

Neither randomization nor the blinding method was used for the selection of animals (8–16-weeks-old). All the experimental protocols involving animals were approved and performed in accordance with the guidelines of the Animal Care and Experimentation Committee of Gunma University (Approval No. 22-076). C57BL/6NCrSlc and hybrid B6D2F1 mice were purchased from the Japan SLC.

### Preparation of oocytes

GV oocytes were collected from the ovaries of female mice, 46 h after the injection of pregnant mare serum gonadotropin or anti-inhibin antibody, CARD HyperOva (Kyudo, Kumamoto, Japan). Antral follicles were suspended in an FHM medium (Sigma, MO, USA). MII oocytes were collected from the oviducts of female mice, 13 h after the injection of CARD HyperOva and 7.5 IU human chorionic gonadotrophin, hCG (ASKA Pharmaceutical, Tokyo, Japan) 48 h apart.

### mRNA synthesis

For the synthesis of the mRNA encoding EGFP, a template cDNA was amplified by a PCR using a T7 promoter-tagged primer (5’- GGGTAATACGACTCACTATAGGGGCCACCATGGTGAGCAAGGGCGAGATACGACTCACTATAGGGGCAACGTGCTGGTTGTTGTGC-3’), reverse primer 5’- GCCACACCAGCCACCACCTTCTG-3’), and high fidelity Taq polymerase, KOD Fx Neo (TOYOBO, Osaka, Japan). RNA synthesis, poly-A tail addition, and further preparation for the injection were performed as previously described (Kato et al., 2016). For the synthesis of mRNA encoding histone H2B-EGFP, a template cDNA with poly-A tail was excised by XhoI digestion of the pcDNA3.1-histone H2B-EGFP-polyA83 plasmid (Hatano et al., 2022; Yamagata et al., 2005). RNA synthesis was carried out using the mMESSAGE-mMACHINE T7 ultra kit (Thermo Fisher Scientific Inc., MA, USA).

### Electroporation

EP was performed using the electroporator NEPA21 (NEPA GENE., Chiba, Japan), the same as the TAKE method (Kaneko and Mashimo, 2015). Up to 40 oocytes or follicles were placed in line between 5-mm gap electrodes on a glass slide CUY520P5 (NEPA GENE) that was filled with 40 μL of Opti-MEM (Thermo Fisher Scientific Inc.) containing 200 μg/mL of mRNA. The oocytes or follicles were then electroporated with a poring pulse (voltage: 225 V; pulse width: 2.0 ms; pulse intervals: 50 ms; number of pulses: 4; decay percentage: 10%; polarity of electrodes: plus) followed by transfer pulses (voltage: 20 V; pulse width: 50 ms; pulse intervals: 50 ms; number of pulses: 5; decay percentage: 40%; polarity: plus/minus). The electroporated oocytes were then cultured in a 20-μL droplet of Opti-MEM for 3 mins followed by incubation in KSOM (Kyudo, Kumamoto, Japan) at 37 °C under 5% CO_2_ in the air. For MII oocytes, survival was determined by morphological observation after 30 mins in the KSOM incubation. The electroporated GV oocytes were immediately subjected to in vitro maturation as described below, and survival was determined by morphological observation of the cumulus-denuded oocytes after the IVM and in vitro fertilization (IVF), approximately 19 h after the electroporation.

### Microinjection into the GV oocytes

Microinjection of mRNA into the GV oocytes was performed according to previous studies (Kato et al., 2016; Umeda et al., 2020). Briefly, antral follicles were suspended in an FHM medium supplemented with 250 μM of dibutyryl-cyclic AMP (dbcAMP), and punctured with 26 G needles (Terumo, Tokyo, Japan). After dissociating the cumulus cells by pipetting, the oocytes were transferred to a medium supplemented with 250 μM of dbcAMP and 5 μg/ml of CCB (Sigma); thereafter the injections of mRNAs (200 ng/μL) into the oocytes were performed using a piezo-micromanipulator with a glass capillary needle. The oocytes were washed three times and then presented for IVM.

### In vitro maturation and fertilization

For the C57BL/6NCrSlc GV oocytes introduced with EGFP mRNA, the GV oocytes were washed twice in Opti-MEM drops following EP. Thereafter, maturation was induced by incubating them in a TYH medium (Toyoda, 1971) containing 10% (vol/vol) fetal bovine serum, FBS (Thermo Fisher Scientific Inc.) for 17 h at 37 °C under 5% CO_2_ in the air as previously described (Kato et al., 2016; Umeda et al., 2020). For the B6D2F1 GV oocytes introduced with histone H2B-EGFP mRNA that were presented for further developmental ability tests or live-cell imaging, the GV oocytes were washed twice in Opti-MEM drops following EP, and maturation was induced by incubation in alpha-MEM (Thermo Fisher Scientific Inc.) containing 5% (vol/vol) FBS, 0.1 of IU/mL follicle stimulating hormone (MSD, Tokyo, Japan), 1.2 IU/mL of hCG (ASKA Pharmaceutical), and 4 ng/mL of epidermal growth factor (Thermo Fisher Scientific Inc.) for 17 h at 37 °C under 5% CO_2_ in the air (Morohaku et al., 2016). For IVF, spermatozoa from a B6D2F1 epididymis were pre-incubated in an mHTF medium (Kyudo) and inseminated at a sperm concentration of 1.5×10^5^ per mL for 1.5 h. Excess spermatozoa were washed out and the oocytes were incubated in the KSOM medium (Kyudo) for further development.

### Confocal fluorescent imaging

Low-invasive fluorescent imaging of the embryos was performed as previously described (Morita et al., 2021; Yamagata et al., 2009). Briefly, confocal images at 41 z-axis planes with 2 μm increments for a 488 nm laser were obtained every 30 min using a confocal scanner unit, CV1000 (Yokogawa Electric Corporation, Tokyo, Japan). The fluorescent images for the evaluation of the introduced EGFP mRNA and the images of the pronuclear (PN) stage embryos after histone H2B-EGFP expression were also acquired on the CV1000. The images of the blastocyst embryos after in vitro culture were acquired on an inverted microscope IX-71 (Olympus, Tokyo, Japan) equipped with a confocal scanner unit W-1(Yokogawa Electric Corporation).

### Statistics and reproducibility

Experiments were repeated at least three times for each batch. Statistical analyses were performed using Graph-Pad Prism 8 software (GraphPad Software, MA, USA). Sample sizes were determined based on previous and similar reports. Morphological abnormality of oocytes were examined at every collection of them from vivo and abnormal oocytes were excluded. The tests were not blinded nor randomized because of shortage of expertized members to execute experiments. Data were checked for the equality of variance using F-test. The means among more groups were examined using a one-way analysis of variance (ANOVA) test. The bars represent mean ± SEM. Statistical significance was set as P-value < 0.05.

### Data and materials availability statement

Data and materials supporting the findings of this manuscript are available from the corresponding authors upon reasonable request. All relevant data can be found within the article and its supplementary information.

## Acknowledgements

A plasmid, pcDNA3.1-histone H2B-EGFP-polyA83, was kindly provided by Professor Kazuo Yamagata of Kindai University, Japan. We also would like to thank Editage (www.editage.jp) for English language editing.

## Author contributions

Y.S. and K.Sato established the essential research concepts for this study. Y.S. and E.S. performed the assays using mouse oocytes/embryos. K. Sasaki established and directed IVM experiments. Y. S. and K. Sato mainly prepared the manuscript.

## Conflict of interest

No conflict of interest declared.

## Funding

This study was supported by the Japan Society for the Promotion of Science KAKENHI (Grant Number 19H05711 and 20H00466 to K. Sato, and 21K15000 to K. Sasaki), and Mochida Memorial Foundation for Medical and Pharmaceutical Research, Takeda Science Foundation (Grant Number 2022093867), and The Cell Science Research Foundation to Y. S.

## Notes

### Competing Interest Statement

The authors have declared no competing interest.

### Summary of Updates

Title have been updated to describe the concept of this article more concise. Summary blurb, running title, data availability, author contribution, and conflict of interest are added to clarify the property of this article.

## References

Bakloushinskaya I (2022) Chromosome changes in soma and germ line: Heritability and evolutionary outcome. Genes (Basel) 13: 602. doi:10.3390/genes13040602

Conti M, Franciosi F (2018) Acquisition of oocyte competence to develop as an embryo: Integrated nuclear and cytoplasmic events. Hum Reprod Update 24: 245–266. doi:10.1093/humupd/dmx040

Gilchrist RB, Smitz J (2023) Oocyte in vitro maturation: Physiological basis and application to clinical practice. Fertil Steril 119: 524–539. doi:10.1016/j.fertnstert.2023.02.010

Hatano Y, Mashiko D, Tokoro M, Yao T, Yamagata K (2022) Chromosome counting in the mouse zygote using low-invasive super-resolution live-cell imaging. Genes Cells 27: 214–228. doi:10.1111/gtc.12925

Hikabe O, Hamazaki N, Nagamatsu G, Obata Y, Hirao Y, Hamada N, Shimamoto S, Imamura T, Nakashima K, Saitou M, et al (2016) Reconstitution in vitro of the entire cycle of the mouse female germ line. Nature 539: 299–303. doi:10.1038/nature20104

Kaneko T, Mashimo T (2015) Simple genome editing of rodent intact embryos by electroporation. PLoS One 10: e0142755. doi:10.1371/journal.pone.0142755

Kaneko T, Sakuma T, Yamamoto T, Mashimo T (2014) Simple knockout by electroporation of engineered endonucleases into intact rat embryos. Sci Rep 4: 6382. doi:10.1038/srep06382

Kato K, Satouh Y, Nishimasu H, Kurabayashi A, Morita J, Fujihara Y, Oji A, Ishitani R, Ikawa M, Nureki O (2016) Structural and functional insights into izumo1 recognition by juno in mammalian fertilization. Nat Commun 7: 12198. doi:10.1038/ncomms12198

Kohama T, Masago M, Tomioka I, Morohaku K (2022) In vitro production of viable eggs from isolated mouse primary follicles by successive culture. J Reprod Dev 68: 38–44. doi:10.1262/jrd.2021-095

Morita A, Satouh Y, Kosako H, Kobayashi H, Iwase A, Sato K (2021) Clathrin-mediated endocytosis is essential for the selective degradation of maternal membrane proteins and preimplantation development. Development 148: dev199461. doi:10.1242/dev.199461

Morohaku K, Hirao Y, Obata Y (2017) Development of fertile mouse oocytes from mitotic germ cells in vitro. Nat Protoc 12: 1817–1829. doi:10.1038/nprot.2017.069

Morohaku K, Tanimoto R, Sasaki K, Kawahara-Miki R, Kono T, Hayashi K, Hirao Y, Obata Y (2016) Complete in vitro generation of fertile oocytes from mouse primordial germ cells. Proc Natl Acad Sci U S A 113: 9021–9026. doi:10.1073/pnas.1603817113

Qin P, Parlak M, Kuscu C, Bandaria J, Mir M, Szlachta K, Singh R, Darzacq X, Yildiz A, Adli M (2017) Live cell imaging of low- and non-repetitive chromosome loci using crispr-cas9. Nat Commun 8: 14725. doi:10.1038/ncomms14725

Sakakibara Y, Hashimoto S, Nakaoka Y, Kouznetsova A, Hoog C, Kitajima TS (2015) Bivalent separation into univalents precedes age-related meiosis i errors in oocytes. Nat Commun 6: 7550. doi:10.1038/ncomms8550

Schroeder AC, Eppig JJ (1984) The developmental capacity of mouse oocytes that matured spontaneously in vitro is normal. Dev Biol 102: 493–497. doi:10.1016/0012-1606(84)90215-x

Secomandi L, Borghesan M, Velarde M, Demaria M (2022) The role of cellular senescence in female reproductive aging and the potential for senotherapeutic interventions. Hum Reprod Update 28: 172–189. doi:10.1093/humupd/dmab038

Thomas C, Cavazza T, Schuh M (2021) Aneuploidy in human eggs: Contributions of the meiotic spindle. Biochem Soc Trans 49: 107–118. doi:10.1042/BST20200043

Toyoda Y, Yokoyama, M. & Hoshi, T. (1971) Studies on the fertilization of mouse egg in vitro. Jpn J Anim Reprod 16: 147–151. doi:10.1262/jrd1955.16.152

Umeda R, Satouh Y, Takemoto M, Nakada-Nakura Y, Liu K, Yokoyama T, Shirouzu M, Iwata S, Nomura N, Sato K, et al (2020) Structural insights into tetraspanin cd9 function. Nat Commun 11: 1606. doi:10.1038/s41467-020-15459-7

Vanderhyden BC, Armstrong DT (1989) Role of cumulus cells and serum on the in vitro maturation, fertilization, and subsequent development of rat oocytes. Biol Reprod 40: 720–728. doi:10.1095/biolreprod40.4.720

Vazquez-Diez C, FitzHarris G (2018) Causes and consequences of chromosome segregation error in preimplantation embryos. Reproduction 155: R63–R76. doi:10.1530/REP-17-0569

Yamagata K, Suetsugu R, Wakayama T (2009) Long-term, six-dimensional live-cell imaging for the mouse preimplantation embryo that does not affect full-term development. J Reprod Dev 55: 343–350. doi:10.1262/jrd.20166.

Yamagata K, Ueda J (2013) Long-term live-cell imaging of mammalian preimplantation development and derivation process of pluripotent stem cells from the embryos. Dev Growth Differ 55: 378–389. doi:10.1111/dgd.12048

Yamagata K, Yamazaki T, Yamashita M, Hara Y, Ogonuki N, Ogura A (2005) Noninvasive visualization of molecular events in the mammalian zygote. Genesis 43: 71–79. doi:10.1002/gene.20158

Yamamoto T, Honda S, Ideguchi I, Suematsu M, Ikeda S, Minami N (2023) A more accurate analysis of maternal effect genes by sirna electroporation into mouse oocytes. J Reprod Dev 69: 118–124. doi:10.1262/jrd.2022-122

Yoshimatsu S, Okahara J, Sone T, Takeda Y, Nakamura M, Sasaki E, Kishi N, Shiozawa S, Okano H (2019) Robust and efficient knock-in in embryonic stem cells and early-stage embryos of the common marmoset using the crispr-cas9 system. Sci Rep 9: 1528. doi:10.1038/s41598-018-37990-w

Zielinska AP, Bellou E, Sharma N, Frombach AS, Seres KB, Gruhn JR, Blayney M, Eckel H, Moltrecht R, Elder K, et al (2019) Meiotic kinetochores fragment into multiple lobes upon cohesin loss in aging eggs. Curr Biol 29: 3749–3765 e3747. doi:10.1016/j.cub.2019.09.006

