## Supplemental Figures for "Improved mRNA electroporation into immature oocytes to examine protein dynamics during development"

### Sfig.1

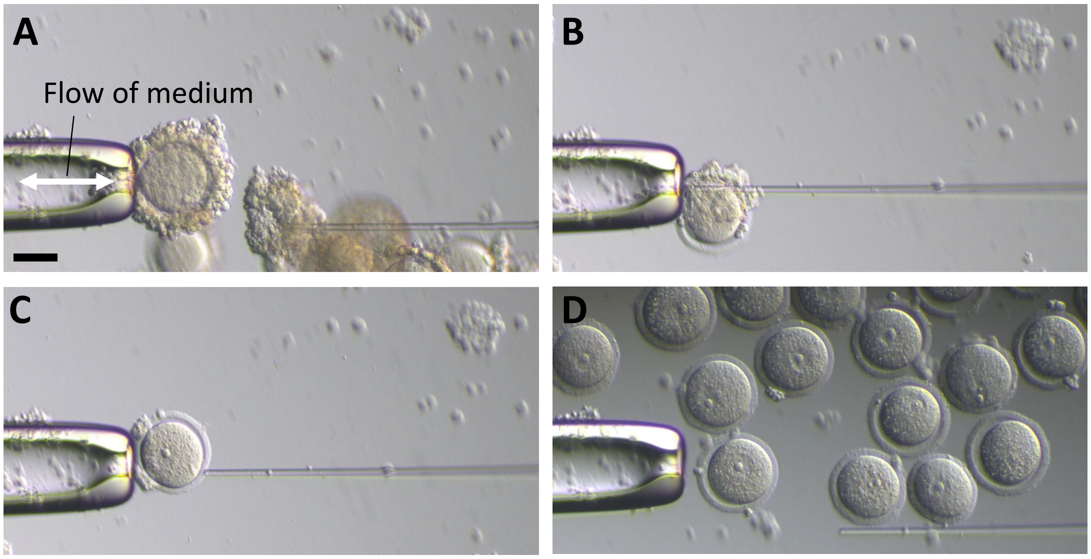


**Supplemental Figure 1. Removal of cumulus cells with a micromanipulator.** A) Aspirated cumulus cells in the holding needle (left side). B and C) Changing an angle of GV oocyte with needles for repetitive aspirations. D) Harvested GV oocytes (cumulus-denuded). Bar: 40 µm.

**Sfig.2**

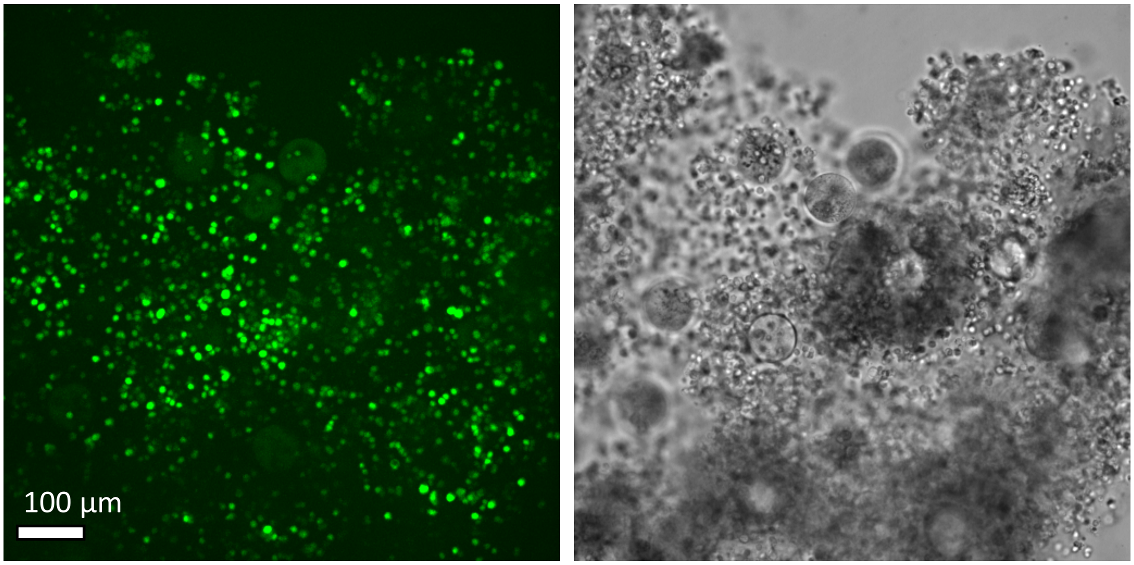


**Supplemental Figure 2. Introduction of mRNA into the cumulus cells.** Confocal fluorescent (max projection of 80 µm slices, in height: left) and bright field images (right) are indicated.

**Sfig.3**

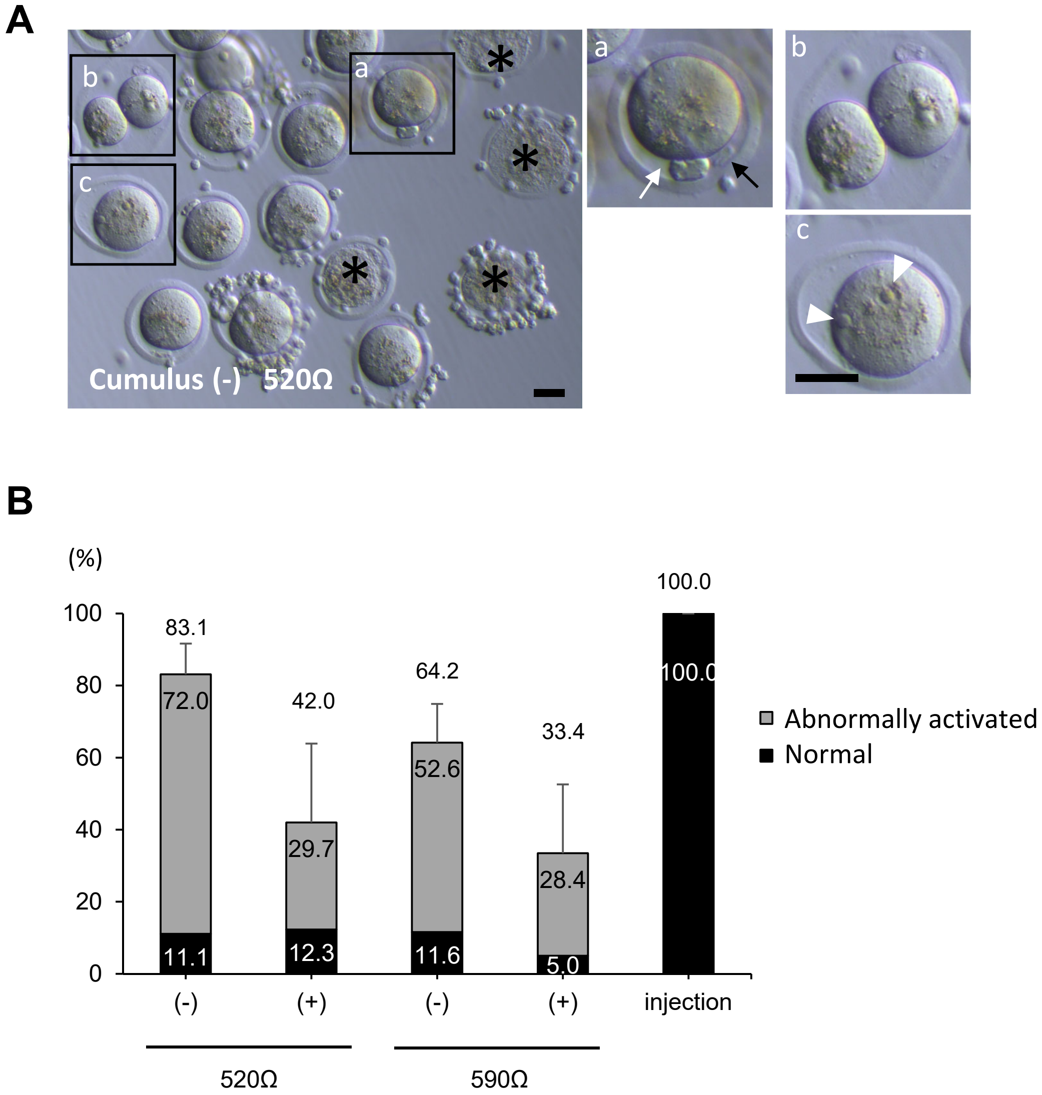


**Supplemental Figure 3. EP into MII oocytes.** A) Abnormally activated oocytes observed after EP to MII oocytes at 520 Ω (cumulus (-)). Examples of oocytes activated without fertilization that spawn not only the first polar body (black arrow) but also the second polar body (white arrow), asymmetrically divided or fragmented oocytes, and oocytes that have PN-like structure(s), are indicated in a, b, and c, respectively. Asterisks indicate dead (ruptured) oocytes. Bars: 40 µm. B) Averaged rates of oocyte survival after 8 h of incubation in the KSOM medium. Percentages of total survival are indicated (including abnormally activated oocytes).

Movie. 1

Histone H2B-EGFP, cumulus (++)

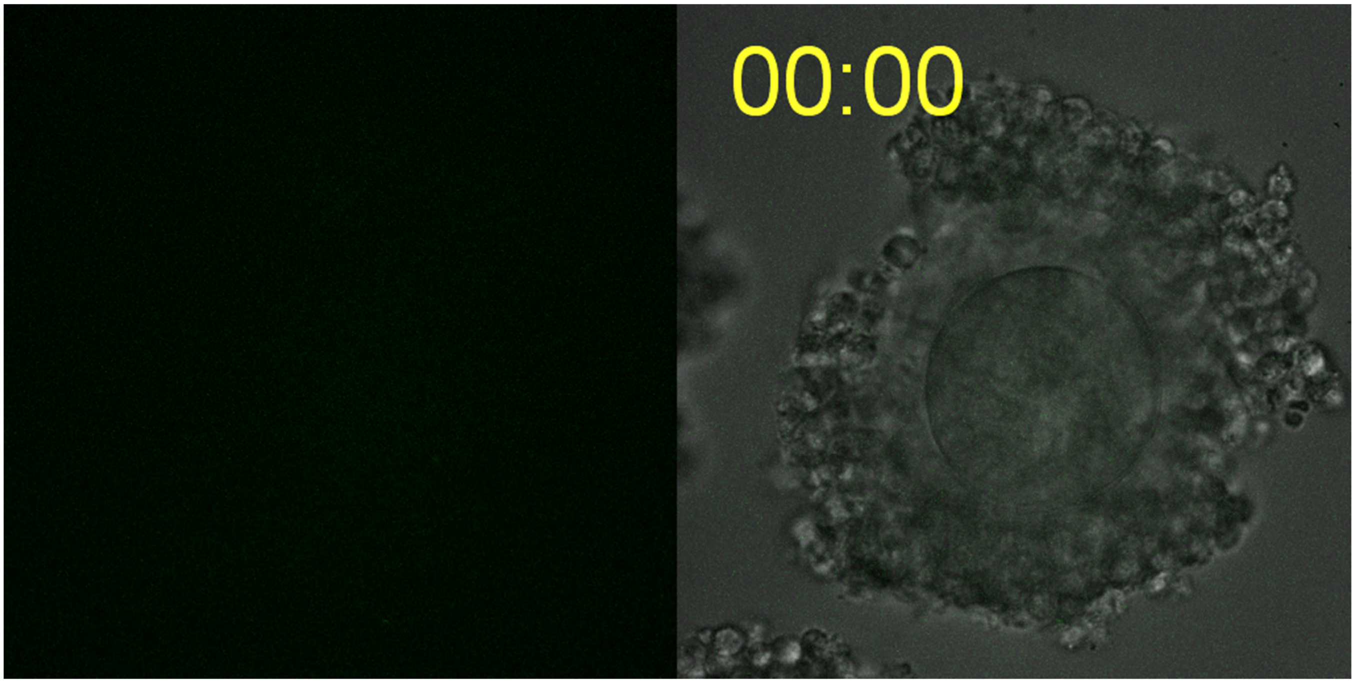


**Supplemental Movie 1** Time-lapse movie of a cumulus (++) GV oocyte after histone H2B-EGFP mRNA EP (maximum intensity projection).

Movie. 2

Histone H2B-EGFP, cumulus (-)

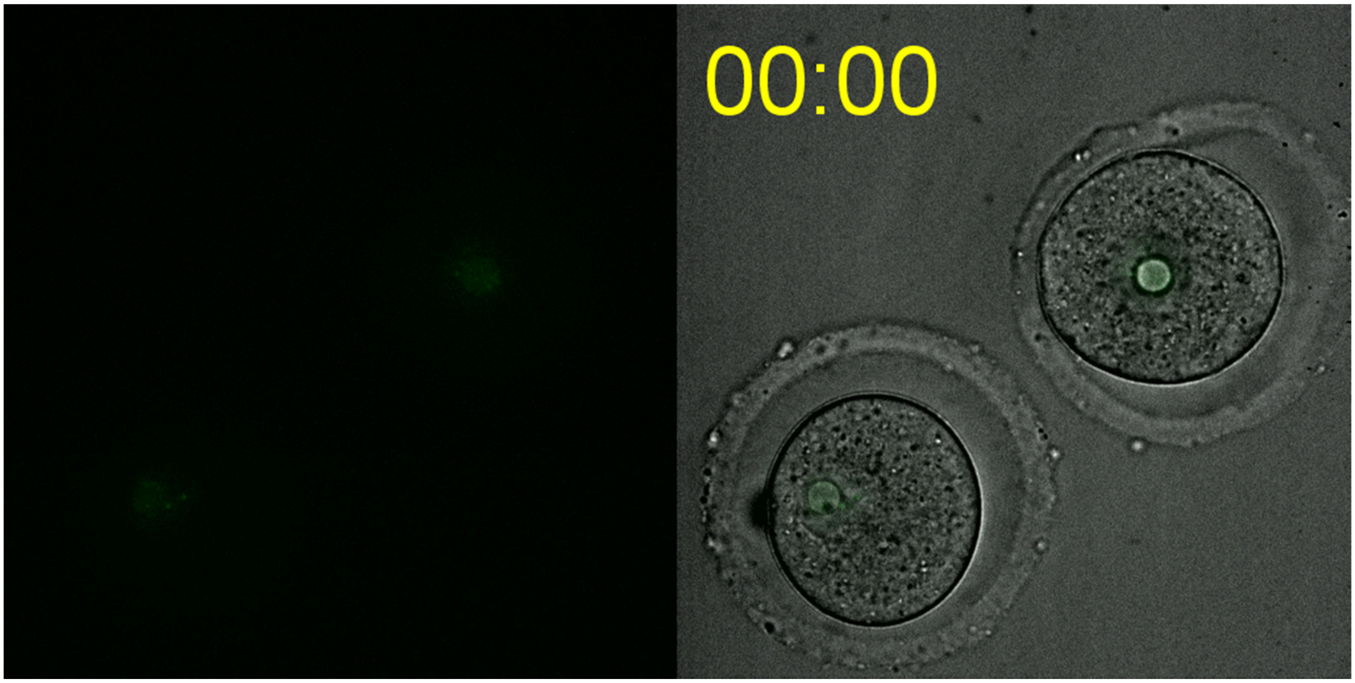


**Supplemental Movie 2** Time-lapse movie of a cumulus (-) GV oocyte after histone H2B-EGFP mRNA EP (maximum intensity projection).
